# Bioscience-scale automated detection of figure element reuse

**DOI:** 10.1101/269415

**Authors:** Daniel E. Acuna, Paul S. Brookes, Konrad P. Kording

## Abstract

Scientists reuse figure elements sometimes appropriately, e.g. when comparing methods, and sometimes inappropriately, e.g. when presenting an old experiment as a new control. To understand such reuse, automatically detecting it would be important. Here we present an analysis of figure element reuse on a large dataset comprising 760 thousand open access articles and 2 million figures. Our algorithm detects figure region reuse, while being robust to rotation, cropping, resizing, and contrast changes, and estimates which of the reuses have biological meaning. Then a three-person panel analyzes how problematic these biological reuses are using contextual information such as captions and full texts. Based on the panel reviews, we estimate that 9% of the biological reuses would be unanimously perceived as at least suspicious. We further estimate that 0.6% of all articles would be unanimously perceived as fraudulent, with inappropriate reuses occurring 43% across articles, 28% within article, and 29% within a figure. Our tool rapidly detects image reuse at scale, promising to be useful to a broad range of people that campaign for scientific integrity. We suggest that a great deal of scientific fraud will be, sooner or later, detectable by automatic methods.

## Introduction

A good amount of scientific misconduct has been found through their inappropriate reuse of figure elements. For example, the infamous stimulus-triggered acquisition of pluripotency (STAP) through stress articles were retracted after several of their figures were found to be inappropriately reused (*1*). Similar investigations often appear in the popular news (*2*). Detecting these types of reuses, however, is tedious (*3*). While the US Office of Research Integrity (ORI) shares several tools to aid in such detection (*4*), ORI reports on approximately 10 new cases of scientific misconduct per year but it is unclear how many cases they open and how many of them involve images (*5*). Also, ORI does not proactively review potential fraud unless it is reported by anonymous sources. Automatically detecting figure reuse could make the process of fraud fighting more efficient.

There have been several proposals to understand the extent of the problem of image reuse in the biological sciences. Recently, Bik, Casadevall and Fang (*6*) manually examined several thousand articles and their images to find problematic reuses. They found that 1.9% of articles analyzed had some deliberate manipulation. Similarly, for the detection of other types of problems, such as statistical inconsistencies, there are some works in progress (*7*). While there are some prototypes to scale the process (e.g., (*8*, *9*)), their effectiveness has not been widely tested. In general, these methods require significant effort and they only work within a figure of a single article. Methods that automatically scale across figures and across articles are lacking.

Here we analyzed all figures published in the PubMed Open Access Subset (PMOS) dataset by 2015, containing more than 2 million figures. We first develop a pipeline for automatic detection of figure element reuse across sets of papers published by the same first or last author. We then develop a protocol to let a set of reviewers manually check the detected image element reuses. Overall, we found that our method is effective in detecting large-scale potentially problematic instances of reuse.

## Materials and methods

### Materials

We analyzed 760,036 articles from the PMOS repository obtained in early 2015. This repository provides the full text of the articles, PDFs, images and datasets associated with each. There are 2,628,959 images contained in these articles.

#### Data clean up

Not all images represent figures of the articles. Many of them represent equations contained within the articles and therefore we removed them from the analysis. Each article contained an average of 4.78 images (SD 3.79). There are 4,324 journals in this dataset, with the biggest 5 journals being PLoS ONE, The Journal of Cell Biology, The Journal of Experimental Medicine, Acta Crystallographica Section E: Structure Reports Online, and the British Journal of Cancer. However, the biggest contributors of images are PLOS ONE (23%), Acta Crystallographica Section E: Structure Reports Online (2.4%), The Journal of Cell Biology (2.3%), Nucleic Acids Research (2.2%), and Sensors (Basel, Switzerland) (1.7%).

**Fig. 1.**
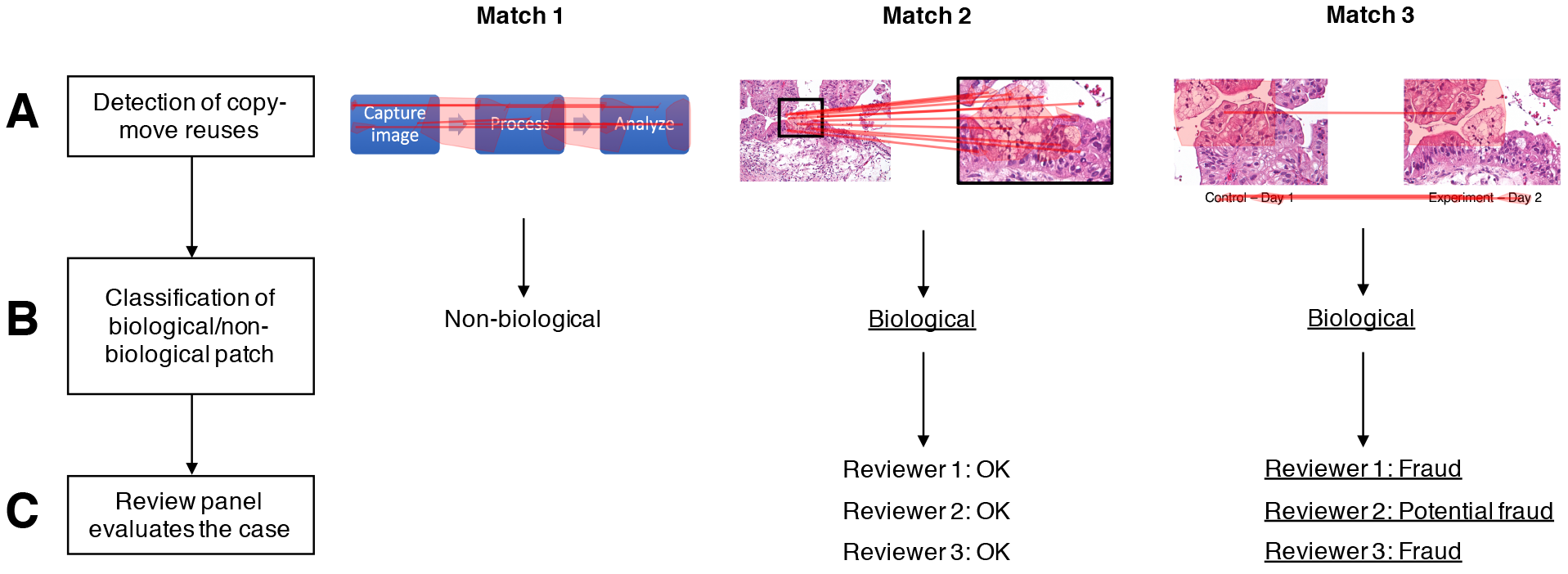
Pipeline for detection and judgement of figure element reuse. **A.** A detection of copy-move reuses is run across all figures. **B.** A pre-trained classifier detects whether the type of copy is biological or not. **C.** A panel of three reviewers goes through each of the copy-move biological copies and tags it as OK, suspicious, potential fraud, or fraud. None of the figure elements shown here are actually fraudulent, these are just schematics.

### Methods

The methods used in this article are a combination of computer assisted tagging of potential image copy-move detection, algorithmic classification of image patches, a combination of human reviewers of copy-move biological patches, and an analysis of the interrater reliability. We will examine each of these results in turn.

**Fig. 2.**
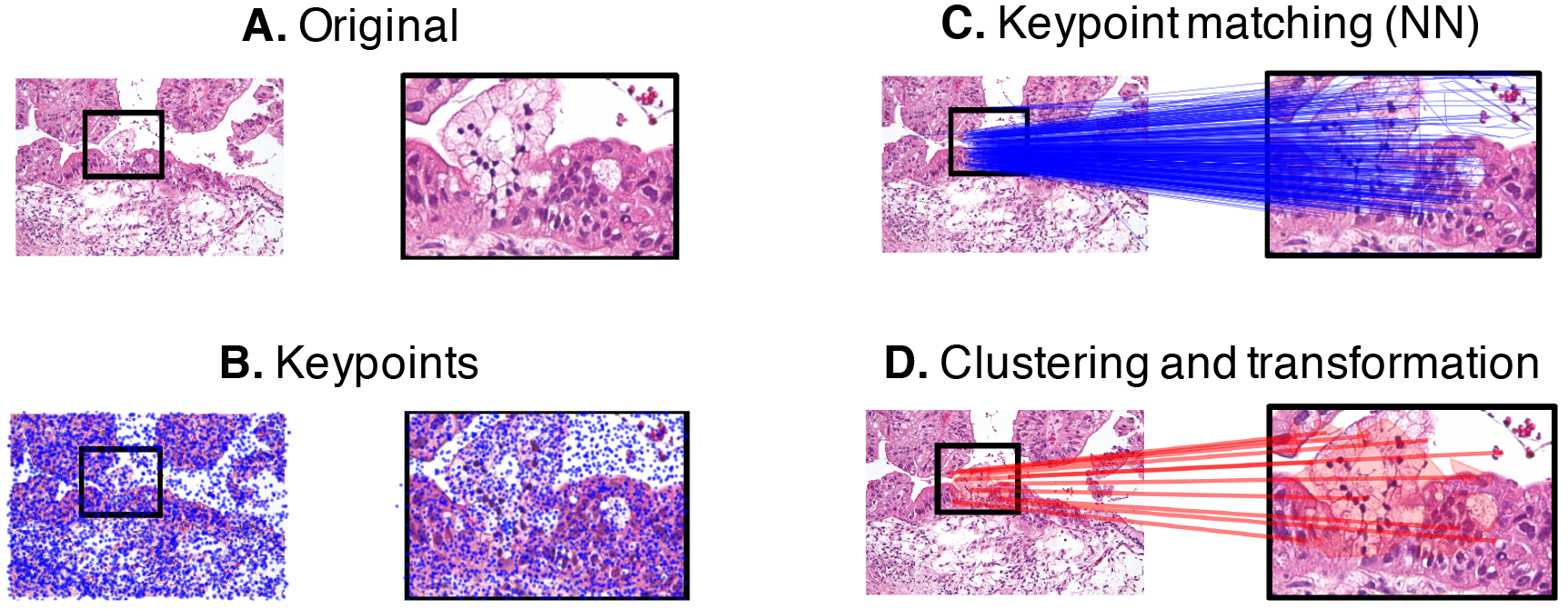
Detection of copy-move reuses. **A.** Original example of an cancerous cell and zoomed portion. **B.** Computation of keypoints (areas of high entropy) **C.** Nearest neighbor matching. **D.** Clustering keypoints, matching across cluster, and affine transform.

#### Copy-move detection algorithm

(Fig. 1A) Algorithmic detection of copy-move elements. The first step to detect any type of reuse is to detect them at large scale. We will first apply the a key-point based method described in (*10*). Specifically, we run our model through the following pipeline:

1. (Fig. 2A) Compute keypoints using SIFT keypoint detection algorithm with a low threshold (*11*). We use this feature since it has a good compromise between speed and robustness (*10*).
2. (Fig 2B) Find a set of matching features using the following procedure: find the two nearest neighbors in Euclidian space across all keypoints detected in the image, if the distance of the nearest neighbor is 60% or less than the distance of the second nearest neighbor. Remove keypoints that do not have matches or that have matches that are less than 40 pixels apart.
3. (Fig. 2C) Perform an agglomerative clustering of keypoints using a minimum distance of 30 pixels to form flat clusters and using a single linkage method. Remove clusters with less than 40×40 square pixels of area.
4. If within a cluster, more than three keypoints are matched against the same cluster, then define those two clusters as matched clusters
5. (Fig. 2D) Use the RanSac algorithm to find the affine transformations between matched clusters. We follow the guidelines of the method described in (*10*). If less than 80% of the keypoints are used by the RanSac algorithm or the mean squared error of the affine transformation is more than 40, we will remove the matched cluster. Also, if the sheer of the estimated transformation is more than 15 degrees, remove the matched cluster.

We focused our analysis on figures from the same last authors and first authors. This was because the time complexity of performing a nearest neighbor search across several million of images is high. Therefore, this analysis leaves outside potential reuses between arbitrary set of authors.

#### Biomedical patches detector

(Fig 1B) Many images in scientific articles share areas that are naturally similar. For example, many images use similar text and shapes to describe areas of a graph, such as axis labels and arrows. Therefore, many of the matches found by the initial phase of the algorithm are these types of copies. We developed an additional step where one of the authors labeled patches detected as copy move elements. This was an active learning approach in which we ask the human to label patches in which a Random Forest algorithm would be most confused about whether the image patch is a Biomedical patch. We sample a set of 20K matches from the copy-move detection previous part and ask the Random Forest to predict the probability that the matches were biomedical matches. We then ranked those matches by the entropy of the prediction and show to the user the matches with highest entropy (prediction closest to 50%). After the human label, we added the data point to the training set and repeated the procedure. For each pair of patches in a matched cluster, the following features are computed:

- color or black and white
- 15-bin three channel histogram of pixel intensity
- From the gray level co-occurrence matrix using 20 pixel distances at 0 angles with 256 levels, symmetric, and normed, extract the following texture features (*12*):

- Expected absolute difference in gray levels
- Expected correlation between gray levels
- 10-bin histogram of gradients with 8 orientations and 16 pixels per cell

The final algorithm for constructing the patch-non-patch classifier is based on Gradient Boosting (GB) with scikit-learn. GB achieved a cross validated Area Under the Curve (AUC) of
865. To present cases to the reviewers that likely involved only biomedical images, we set the classification for the algorithm so that it achieved a 0.5% False Positive Rate. The True Positive Rate of such threshold was 13.1%. This generated a total of 30,870 matches from 17,069 figures from 11,814 articles.

**Fig. 3.**
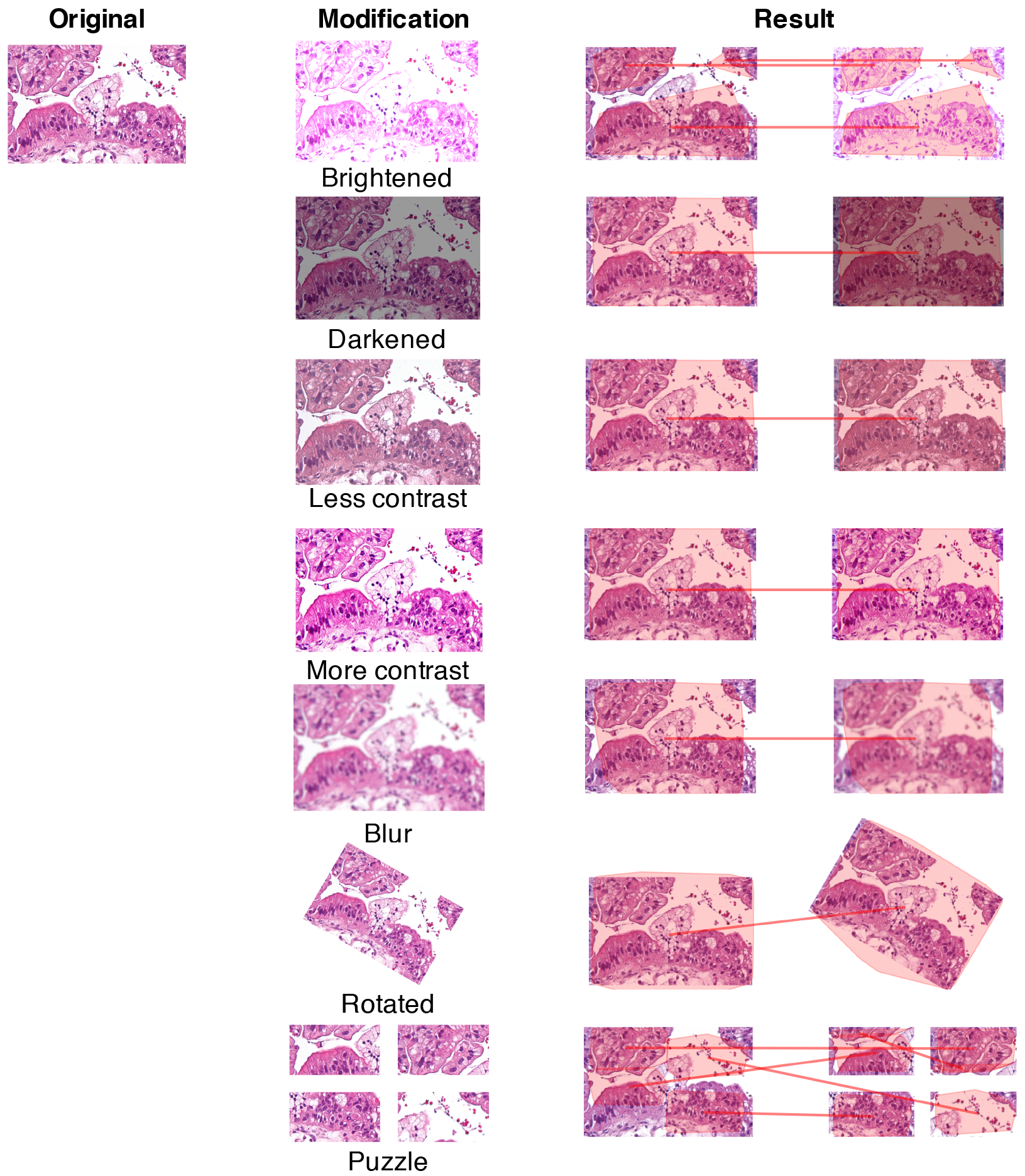
Different types of modifications that the algorithm is able to detect.

#### Human review of potential problematic copies

(Fig 1C) The previous step developed a method to detect copy-move patches that contain biomedical images. The goal of this work, however, is to evaluate the extent of problematic copies. To achieve this, we ran a classification algorithm to detect biological matches, described above. We use a threshold that achieves a 0.5% false positive rate. The idea behind such low false positive was to present human judges with cases where the algorithm is highly confident that there is a Biological sciences image.

We provided a panel of three human reviewers (all authors) with a web-based tool. The web-based tool presented 10,000 cases of potentially problematic cases, which were independently reviewed by the authors of this study. The web-page presented the matches with a link to the PUBMED figure where the judges could review the captions of the figures, thearticles themselves, and elucidate whether the match originated from an inappropriate reuse of a biological image. This secondary evaluation included the reviewers reading figure legends, for example to ascertain whether the same images had been used to represent different experimental conditions.

**Fig. 4.**
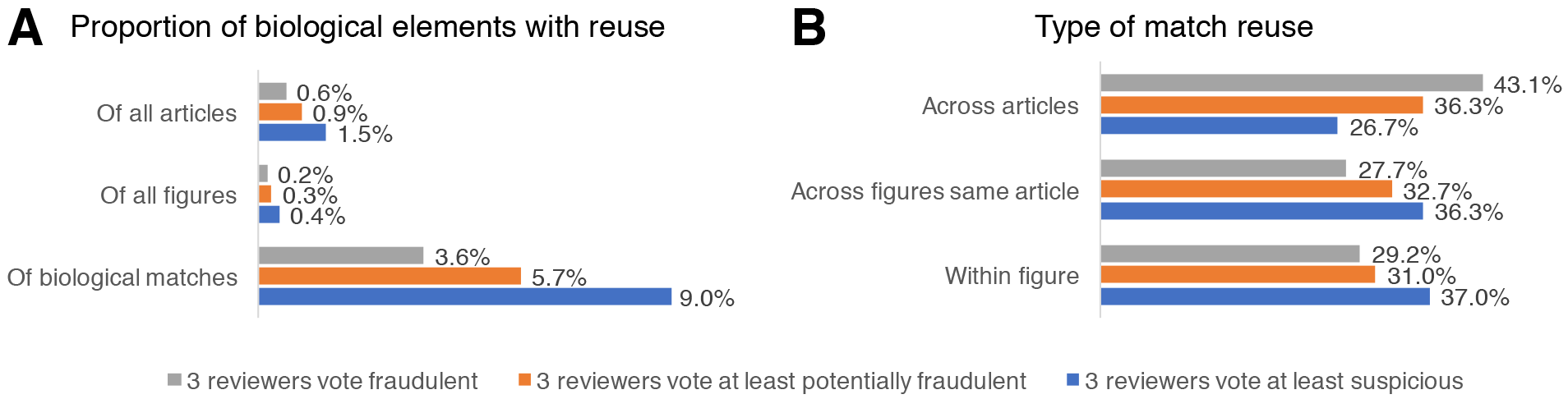
Results of the algorithm and panel review **A.** Of the proportion of biological elements found by the algorithm, a set of reviewers found that at least 0,59% of all articles have fraud. **B.** Given the type of votes for a match, what is the distribution of types of matches that occur. Of the type of matches found, the unanimously fraudulent, 43.1% are matches across articles.

## Results

Here we built a pipeline to automatically detect candidates for inappropriate image reuse. We first removed images that were likely just text or equations represented as images. This left us with around 2 million images. We found on average around IK high-entropy key points per image. This produced a large similarity detection problem, which we solve using approximate nearest neighbor algorithms. We then use a machine learning algorithm to estimate if a patch shows a biological image. Lastly, we use human evaluations of inappropriate reuse. This three-stage pipeline allows us to detect inappropriate image reuse at scale.

There are many kinds of reuses, some of which are necessary. For example, authors usually repeat symbols and axis names within panels in figures. Also, they copy panels between figures of the same paper to make it easier for the users to compare results. Also, authors reuse figures across papers and cite the reused papers accordingly. We considered these cases non-biological because they do not relate to biological phenomena. There are other kinds of biological images that naturally contain many harmless reuses, such as tissue staining. Out of these uninteresting cases, we found that around 6.03% of them are interesting biological reuses. There are many kinds of reuses but only the biological ones were used for our pipeline.

The panel of reviewers was then fed the results of the biologically interesting matches. The three authors acted as reviewers. The reviewers could mark the potential matches as: No problem, suspicious, potential fraud, and likely fraudulent. Additionally, the reviewers could add comments to the annotations. Author 1 went through 3,495 matches, author 2 went through 1,147 matches, and author 3 went through 3,375 matches. Of them, 1,055 matches where reviewed by the three reviewers, 2,193 matches were reviewed by two reviewers, and 466 matches were reviewed by one reviewer. For these 1,055 matches, we found that reviewers agree 88% of the time on whether the matches are at least suspicious, 90% of the time on whether the matches are at least potentially fraudulent, and 94% of the time on whether the matches are likely fraudulent. We additionally found that consistency and agreement of the score is high (ICC(1) = 0.481 [0.445, 0.516], *F*(1054, 2110) = 3.78, *p* < 0.001) when considering matches as random effects, and slightly more consistent when considering matches and reviewers as random effects (ICC(1) = 0.49, *F*(1054, 2108) = 3.89, *p* < 0.001). According to (*13*), an ICC between 0.4 and 0.59 is considered “fair”. On average, reviewers found that 9% of the matches were at least suspicious (author 1: 9.1% (out of 3,495), author 2: 3.6% (out of 1,147), author 3: 10.7% (out of 3,375)), 5.7% at least potentially fraudulent, and 3.6% fraudulent.

Based on the panels’ judgements of the biological matches and the true positive rate of the classified that selected them, we can estimate how many figures and articles are expected to be problematic. Overall, of the articles in PMOS, 1.47% of the articles would be considered suspicious by at least one of a panel of three reviewers, which somewhat resembles the findings in (*6*). 0.59% of articles would be unanimously considered fraudulent (Fig. 4A).

Finally, we examined how matching occurred within articles and across articles. We found that most of the suspicious behavior was found within an article (Fig. 4B), while for cases that were unanimously considered fraudulent, they were more likely to be found across articles (43.1) than when they are considered suspicious (26.7%). Overall, this indicates that almost half of the reuses are happening across articles.

## Conclusion

In this article, we have built a pipeline to analyze potential inappropriate reuse of figures in the biological sciences literature. The analysis relies on a copy-move detection algorithm and then a classification of the potential matches into biological or non-biological. After that, a panel of reviewers reviews the context in which those matches occur. Overall, our results suggest that around 0.59% of the articles in PubMed Open Access would be unanimously considered fraudulent by a panel of three scientists. We now discuss some problems with our analysis.

One potential shortcoming of our results is that some degree of understanding of the intent of the authors behind the reuses is required to draw firm conclusions regarding the benign/fraudulent nature of each case. We may not be familiar with all the techniques presented in the figures. One of the authors, however, has investigated these types of cases extensively (*14*), and the authors of (*6*) have also spoken about the need for a “trained eye” in such matters. It is also possible that the reuses marked as fraudulent are indeed plausible errors during the production of the articles, such as cases from the past (*15*). Standards for scientific image preparation are constantly evolving, and practices that may have been permitted in the past (e.g. undisclosed splicing of western blots) are no longer considered benign. Our plan is to let academic research integrity offices review these types of cases with the identifiable information of the authors but have clear disclosure of the limitations of our method.

Another potential problem is that we are only accessing Open Access publications added to the PubMed repositories and are only reviewing potential reuses within the first and last author’s articles. The dataset analyzed here contains 4,085 Open Access journals out of the estimated 9,455 journals in Open Access (from the Directory of Open Access Journals, doaj.org) and only 760,036 articles of the estimated total of 120 million articles (*16*). However, we focus our analysis on Biomedical sciences which has a much-reduced number of publications. Whether our findings are generalizable to the largely pay-walled biomedical sciences literature at large, remains to be seen. Currently however, widespread access to figures of such articles is not easy, and may lead to interesting ethical problems. A clear conflict of interest also exists, wherein journal publishers may be disinclined to allow access a repository of articles, by researchers whose main goal is to identify problems that arguably should have been detected in the initial peer review process. Also, because we only analyze first and last author’s papers, we are missing on reuses across authors. Extending our approach to all of science and across authors seem highly desirable.

Our method produces findings at a much larger scale than before. We are able to do the bulk of our analysis automatically and let reviewers analyze those cases that are almost certainly biological matches. Additionally, our method can compare reuses across articles, which is very difficult to do if the reviewers of the copies are humans. Even though our method is now trained to detect biomedical images, it would be relatively easy to train it to detect other reuses of artifacts such as graphs or tables. We believe that the widespread adoption of this method by scientific publishers as a screening tool during article submission could act as a deterrent mechanism. We also believe that are many other uses of this technology, such as finding where images and other scientific artifacts are reused in other publications for credit purposes.

Regarding the downstream consequences of the specific cases of image reuse identified herein, an automated tool to notify journal editors, authors, academic research integrity officers and other interested parties is not widely available. Thus, our tool automates what is currently only the first step in a long and laborious process of actually correcting the scientific literature. A potentially serious issue arising from these results is that they may cause harm to careers, and as such we have chosen not to release the results of our automated screening. In the future, as we scale this approach further, we plan on allowing access to the list of papers to institutions that go through an access agreement to protect authors during potential investigations. Overall, the reasons why scientists may commit misconduct are still poorly understood (*17–19*), but regardless of motive the requirement to “tread carefully” when handling such matters remains paramount.

## Acknowledgements

Daniel E. Acuna was generously supported by NSF #1646763, and Daniel E. Acuna and Konrad Kording were supported by PHS Grant P01-NS44393, and John Templeton Foundation. We would like to thank Amazon Web Services’ Cloud Credits for Research. We are grateful for the feedback of Lauran Qualkenbush from Northwestern University.

## References

1. D. Cyranoski, Cell-induced stress. Nature 511, 140 (2014).

2. J. Glanz, A. Armendariz, in New York Times. (New York, 2017), pp. A1.

3. J. Krueger, Forensic Examination of Questioned Scientific Images. Accountability in Research 9, 105–125 (2002).

4. Office of Research Integrity. (2017).

5. N. Gilbert. (Nature Publishing Group, 2009).

6. E. M. Bik, A. Casadevall, F. C. Fang, The prevalence of inappropriate image duplication in biomedical research publications. mBio 7, e00809–00816 (2016).

7. M. B. Nuijten, C. H. Hartgerink, M. A. van Assen, S. Epskamp, J. M. Wicherts, The prevalence of statistical reporting errors in psychology (1985-2013). Behavior research methods 48, 1205–1226 (2016).

8. L. Koppers, H. Wormer, K. Ickstadt, Towards a Systematic Screening Tool for Quality Assurance and Semiautomatic Fraud Detection for Images in the Life Sciences. Science and Engineering Ethics, 1–16 (2016).

9. A. Abbott, Image search triggers Italian police probe. Nature 504, 18 (2013).

10. V. Christlein, C. Riess, J. Jordan, C. Riess, E. Angelopoulou, An evaluation of popular copy-move forgery detection approaches. IEEE Transactions on information forensics and security 7, 1841–1854 (2012).

11. D. G. Lowe, in Computer vision, 1999. The proceedings of the seventh IEEE international conference on. (Ieee, 1999), vol. 2, pp. 1150–1157.

12. R. M. Haralick, K. Shanmugam, Textural features for image classification. IEEE Transactions on systems, man, and cybernetics 3, 610–621 (1973).

13. D. V. Cicchetti, Guidelines, criteria, and rules of thumb for evaluating normed and standardized assessment instruments in psychology. Psychological assessment 6, 284 (1994).

14. P. S. Brookes, Internet publicity of data problems in the bioscience literature correlates with enhanced corrective action. PeerJ 2, e313 (2014).

15. Correction: Preclinical Assessment of FHIT Gene Replacement Therapy in Human Leukemia Using a Chimeric Adenovirus, Ad5/F35. Clinical Cancer Research, (2016).

16. M. Khabsa, C. L. Giles, The number of scholarly documents on the public web. PloS one 9, e93949 (2014).

17. D. Fanelli, R. Costas, F. C. Fang, A. Casadevall, E. M. Bik, Why Do Scientists Fabricate And Falsify Data? A Matched-Control Analysis Of Papers Containing Problematic Image Duplications. bioRxiv, 126805 (2017).

18. A. Zellmer, Family-friendly science. Science 354, 1070–1070 (2016).

19. K. Powell, Young, talented and fed-up: scientists tell their stories. Nature 538, 446–449 (2016).

